# A quantile integral linear model to quantify genetic effects on phenotypic variability

**DOI:** 10.1101/2021.04.14.439847

**Authors:** Jiacheng Miao, Yupei Lin, Yuchang Wu, Boyan Zheng, Lauren L. Schmitz, Jason M. Fletcher, Qiongshi Lu

## Abstract

Detecting genetic variants associated with the variance of complex traits, i.e. variance quantitative trait loci (vQTL), can provide crucial insights into the interplay between genes and environments and how they jointly shape human phenotypes in the population. We propose a quantile integral linear model (QUAIL) to estimate genetic effects on trait variability. Through extensive simulations and analyses of real data, we demonstrate that QUAIL provides computationally efficient and statistically powerful vQTL mapping that is robust to non-Gaussian phenotypes and confounding effects on phenotypic variability. Applied to UK Biobank (N=375,791), QUAIL identified 11 novel vQTL for body mass index (BMI). Top vQTL findings showed substantial enrichment for interactions with physical activities and sedentary behavior. Further, variance polygenic scores (vPGS) based on QUAIL effect estimates showed superior predictive performance on both population-level and within-individual BMI variability compared to existing approaches. Overall, QUAIL is a unified framework to quantify genetic effects on the phenotypic variability at both single-variant and vPGS levels. It addresses critical limitations in existing approaches and may have broad applications in future gene-environment interaction studies.

## Introduction

Human complex phenotypes are shaped by numerous genetic and environmental factors as well as their interactions^1^. Genome-wide association studies (GWAS) have identified tens of thousands of reproducible genetic associations^2^. However, there has been limited success in detecting interactions between human genetic variants and environmental factors (GxE)^3^, in part due to the polygenic nature of human traits, small effect sizes of GxE interactions, and a high multiple testing burden^4^. An alternative approach is to first quantify the overall genetic propensity in the form of polygenic scores (PGS) for each individual, and then test the interactions between PGS and environmental risk factors^5-9^. Here, PGS is a sum of trait-associated alleles across many genetic loci, typically weighted by marginal effect sizes estimated from a GWAS.

However, genetic variants affect not only the level of traits, but also the variability^10-12^. Since the variance of a quantitative phenotype differs across genotype groups of variants involved in GxE interactions, one can use genetic variants associated with the trait variability (vQTL) to screen for candidate GxE interactions^13-16^. The concept of PGS, which estimates the conditional mean of the phenotype^17^, has also been extended into genome-wide summaries of genetic effects on phenotypic variability (vPGS)^18,19^. These scores, which reflect the genetic contribution to outcome plasticity, have suggested unique genetic contributions orthogonal to that of traditional PGS and have achieved some recent successes in GxE studies^18,20^.

Robust vQTL findings can be used to prioritize candidate variants in GxE analysis. vPGS also has the potential to aggregate information across numerous genetic loci and improve both statistical power and biological interpretability of GxE studies. However, existing statistical methods for vQTL and vPGS have limitations^21,22^. Levene’s test (LT)^23^ and deviation regression model (DRM)^13^ are robust to model misspecification but do not adjust for confounding effects on trait variance^24,25^. Additionally, these methods cannot be applied to continuous predictors (e.g., vPGS) because they require the phenotypic mean or median within each category of the predictor (e.g., genotype groups) as input. Heteroskedastic linear mixed models (HLMM) can adjust for covariates but are sensitive to model misspecification and have type-I error inflation when applied to non-normal phenotypes^14,16,26^. Further, although it is straightforward to calculate vPGS using vQTL effects as variant weights, predictive performance of vPGS has not been properly benchmarked due to a lack of statistical metrics for variance prediction. There is a need for a unified framework that can accurately and robustly quantify the genetic effects on phenotypic variability at the single variant as well as the vPGS level.

In this work, we introduce QUAIL (**qua**ntile **i**ntegral **l**inear model), a quantile regression-based framework to estimate genetic effects on the variance of quantitative traits. Our approach can adjust for confounding effects on both the level and the variance of phenotypic outcomes, can be applied to both categorical and continuous predictors, and does not impose strong assumptions on the distribution of phenotypes. We demonstrate the performance of QUAIL through extensive simulations, vQTL mapping for body mass index (BMI) in UK Biobank, GxE enrichment analysis, and vPGS benchmarking and application.

## Results

### Method overview

The goal of vQTL mapping is to identify single nucleotide polymorphisms (SNPs) showing differential variability of a quantitative trait across genotype groups. If a SNP has substantially different effects on trait values given different environmental exposures, it will be a vQTL without conditioning on the environment (**Figure 1A**). Quantile regression estimates the conditional quantile function of a response variable given predictors^27^. If a SNP *G* is a vQTL for trait *Y*, the conditional quantile function will have different regression slopes (i.e., *β*_*τ*_) for different quantile levels *τ* (**Figure 1B, Supplementary Note**)^28^.

**Figure 1.**
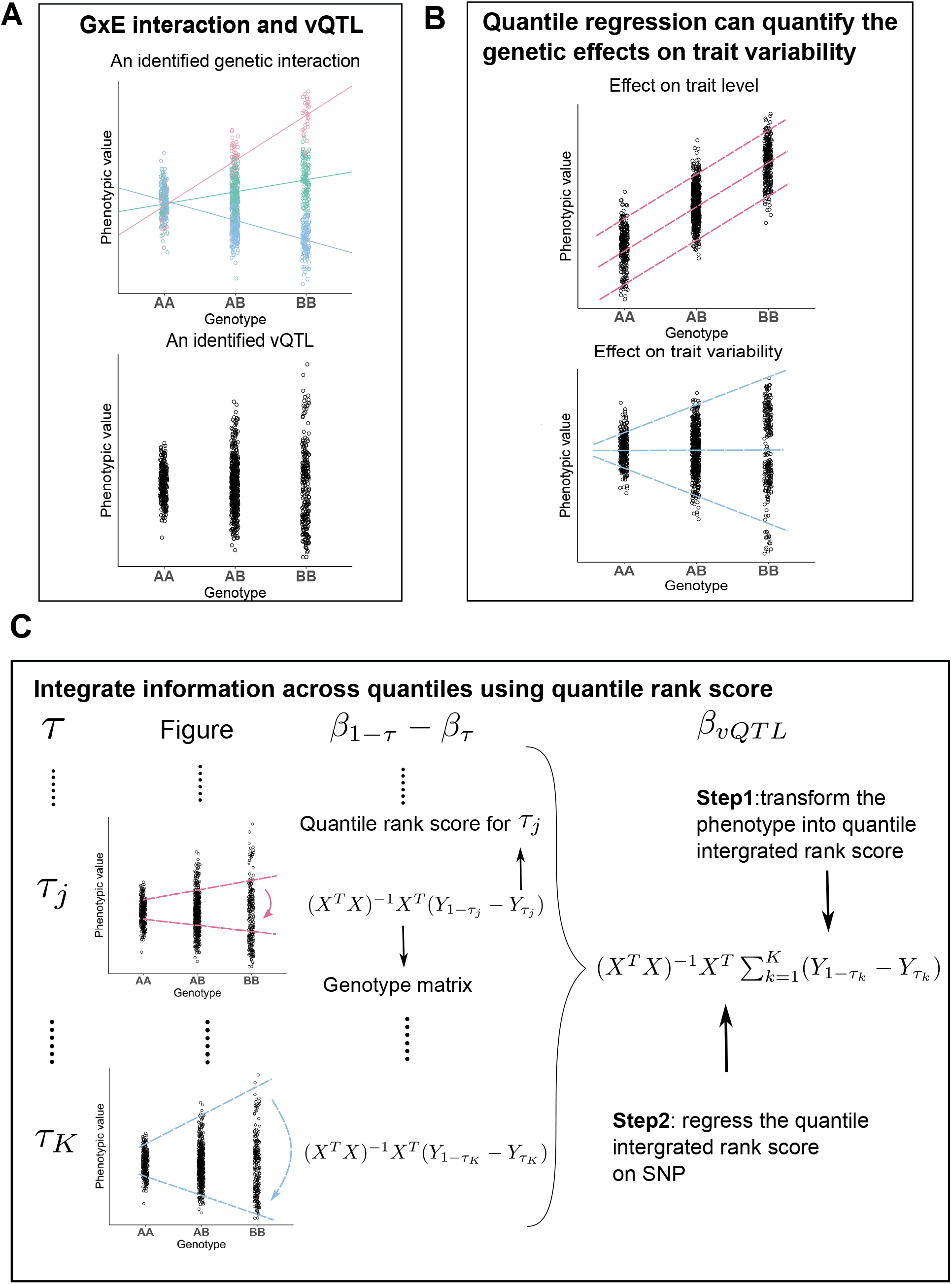
Workflow of QUAIL. **(A)** The phenotypic variance varies across genotype groups in the presence of vQTL and GxE effects. The data points are colored based on the level of the environmental variable. The lines represent genetic effects on the phenotype conditioning on the environmental variable. **(B)** Quantile regression can be used to detect vQTL. The quantile regression slopes will be different across quantile levels if a genetic effect on trait variability exists. **(C)** Workflow of the QUAIL estimation procedure. *τ* indicates a specific quantile level. *β*_1−*τ*_ − *β*_*τ*_ indicates the difference between the regression coefficients of the upper and lower quantile levels. *β*_*vQTL*_ denotes the aggregated genetic effect on trait variability across quantile levels.

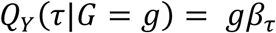

For a pair of quantile levels (1 − *τ, τ*), *τ*∈ (0, 0.5), vQTL effect of a SNP can be quantified using the difference between the regression coefficients of the upper and lower quantile levels, i.e., *β*_1−*τ*_ − *β*_*τ*_.

To aggregate information across all quantile levels and better quantify the vQTL effect on trait *Y*, we introduce a quantile-integrated effect^29^:

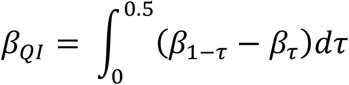

Note that when the SNP is not associated with trait variability, we have *β* _*τ*_ = *β*_1−*τ*_ for any *τ*∈ (0, 0.5). Therefore, testing the vQTL effect of a SNP is equivalent to testing the null *H*_0_: *β*_*QI*_ = 0. However, approximating *β*_*QI*_ using a standard quantile regression fitting procedure involves iterative optimization for numerous quantile levels and thus is computationally challenging in genome-wide analysis. We apply several computational techniques to ensure that QUAIL can efficiently identify vQTL at the genome-wide scale. We first transform the phenotype into an integrated quantile rank score using only trait values and covariates. Next, we regress the transformed phenotype on covariate-adjusted SNP residuals. To estimate integral *β*_*QI*_, QUAIL avoids fitting regressions for a grid of quantile levels. Instead, it only requires fitting two linear regressions per SNP in genome-wide analysis (**Figure 1C**). We present detailed derivations and technical discussions of the QUAIL framework in **Methods** and **Supplementary Note**.

### Simulation results

We performed simulations to compare the empirical performance of QUAIL with four other vQTL methods: DRM^13^, LT^23^, and HLMM with and without inverse normal transformation^16^. We compared the statistical power, false-positive rate (FPR), and ability to adjust for confounding effects for these methods.

We evaluated the FPR and power of all approaches under several simulation scenarios, including 1) three different distributions of the error term to represent various degrees of skewness and kurtosis in the phenotype, and 2) two types of SNP effects on the level and the variance of the phenotype (**Methods**). For FPR simulations, we used a model where the SNP only has effects on the level but not the variance of the phenotype. We used the phenotypic variance explained (PVE) by the SNP to control the magnitude of effects. For power simulations, we simulated quantitative trait values using a GxE interaction model without genetic main effects (**Methods**), where the SNP only has effects on the variance but not the mean of the phenotype. We used PVE by the GxE interaction to control the magnitude of variance effects.

Throughout all simulations, QUAIL maintained well-controlled type-I error regardless of the phenotypic distribution and showed superior power when the phenotype is not normally distributed. When the phenotype follows a normal distribution, all methods control the type-I error well and HLMM is more powerful than other approaches (**Figures 2A** and **2D**). When the phenotype is kurtotic (**Figures 2B** and **2E**) or skewed (**Figures 2C** and **2F**), HLMM shows inflated type-I error. QUAIL, DRM, and LT are robust to the skewness and kurtosis of the phenotype. QUAIL is the most powerful method when the phenotype is kurtotic and shows similar power to DRM and LT when the phenotype is skewed.

**Figure 2.**
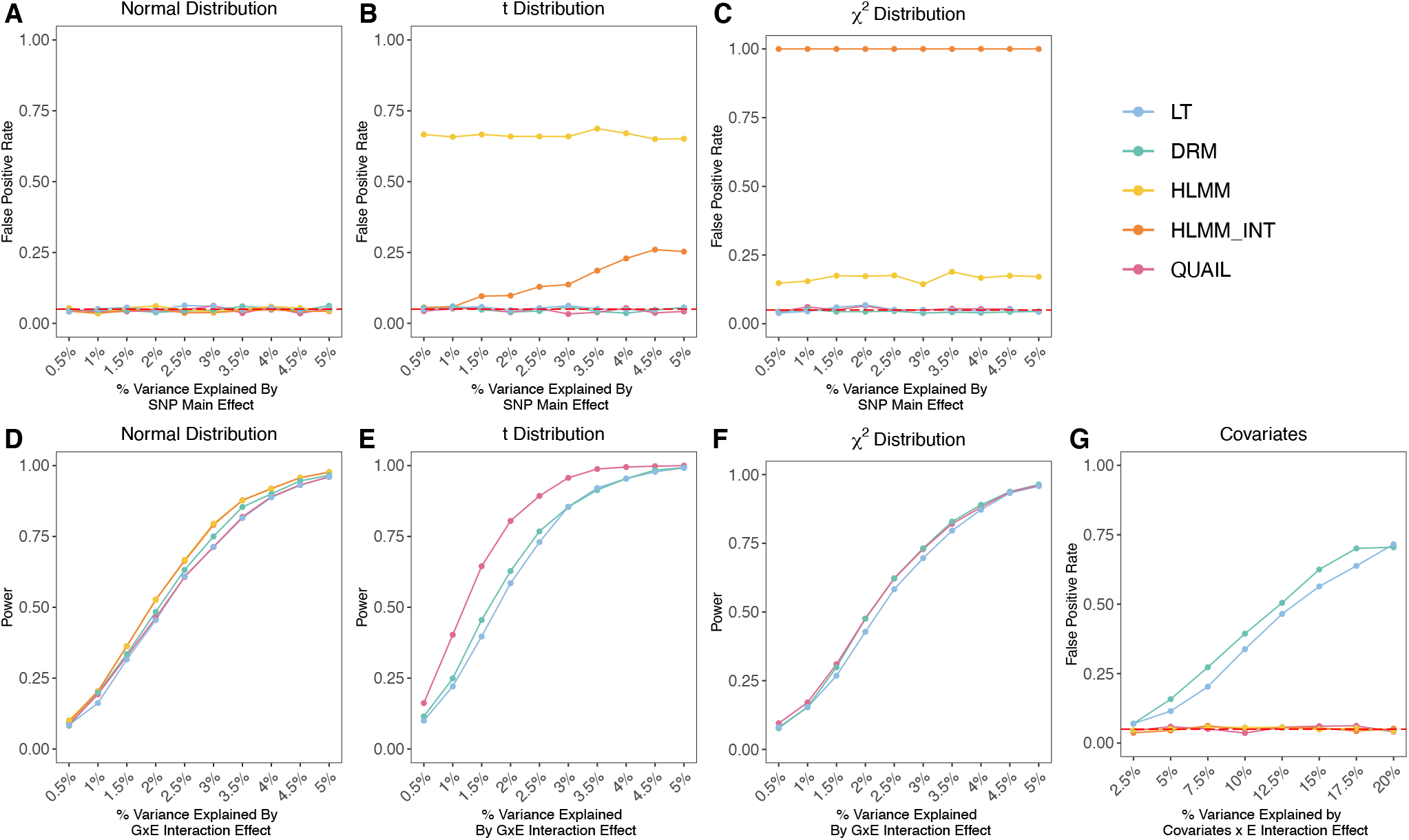
Simulation results. Panels **A-C** compare the false positive rate (FPR), and panels **D-F** compare the statistical power of QUAIL and other vQTL methods under three different phenotypic distributions. **(A, D)** The phenotype follows a normal distribution. **(B, E)** The phenotype follows a t distribution. **(C, F)** The phenotype follows a *χ*^2^ distribution. **(G)** FPR of vQTL methods when confounding effect on trait variability is present.

To examine the ability to adjust for confounding effects on phenotypic variance (**Supplementary Figure 1**), we first simulated a SNP and a correlated covariate. Next, we simulated the phenotype using a covariate x E interaction model (**Methods**). We did not include the SNP in the data-generating model, so the SNP has no causal effect on the phenotype. We also did not include a main effect of the covariate to ensure that the covariate does not affect trait levels. We applied all vQTL methods to test whether the SNP is associated with the variance of the phenotype (**Figure 2G**). QUAIL and HLMM maintained a well-controlled type-I error rate and successfully adjusted for the covariate’s effect on variance. DRM and LT showed inflated FPR, suggesting a lack of robustness to confounding. We summarize the properties of these vQTL methods in **Table 1**.

**Table 1.** A summary of key properties of vQTL methods.

|  | Adjust for covariates' effects<br>on trait level and variability | Robust to non-normal phenotype | Continuous<br>predictor |
| --- | --- | --- | --- |
| QUAIL | Both | Yes | Yes |
| DRM | Only on trait level | Yes | No |
| Levene's test | Only on trait level | Yes | No |
| HLMM | Both | No | Yes |
| HLMM_INT | Both | No | Yes |

### Identifying vQTL for BMI in UK Biobank

We applied QUAIL to perform genome-wide vQTL analysis on unrelated samples of European descent in the UK Biobank. After sample quality control (QC), 375,791 individuals with genotype data and BMI measurements were included in the analysis (**Methods**). We adjusted for sex, age, genotyping array, and genetic principal components (PCs) in the analysis. For comparison, we also applied DRM and HLMM to the same dataset. Both variance effect (HLMM_Var) and dispersion effect (HLMM_Disp) estimates were obtained from HLMM. We applied the inverse normal transformation to BMI before fitting HLMM. LT was omitted in this analysis due to its near identical performance compared to DRM.

**Figure 3A** shows the Manhattan plot for QUAIL vQTL. We identified 49 significant (P< 5.0e-8), approximately independent (pairwise *r*^2^ < 0.01) loci (**Supplementary Table 1**). QUAIL identified more loci than other approaches (**Figure 3B**). Among these 49 loci, 11 are novel vQTL not identified by other approaches. The quantile-quantile plot of QUAIL vQTL hints at inflation (*λ*_*GC*_ = 1.339; **Supplementary Figure 2**), but the intercept of linkage disequilibrium (LD) score regression is 1.003, which suggests polygenic vQTL associations rather than unadjusted confounding. Furthermore, we applied ashR^30^ to estimate the fraction of non-null associations in genome-wide vQTL statistics. We estimated that 85% of all common SNPs have non-zero effects on the variability of BMI, which is consistent with an “omnigenic” model for BMI^6^ and suggests that more loci with small variance effects are yet to be identified.

**Figure 3.**
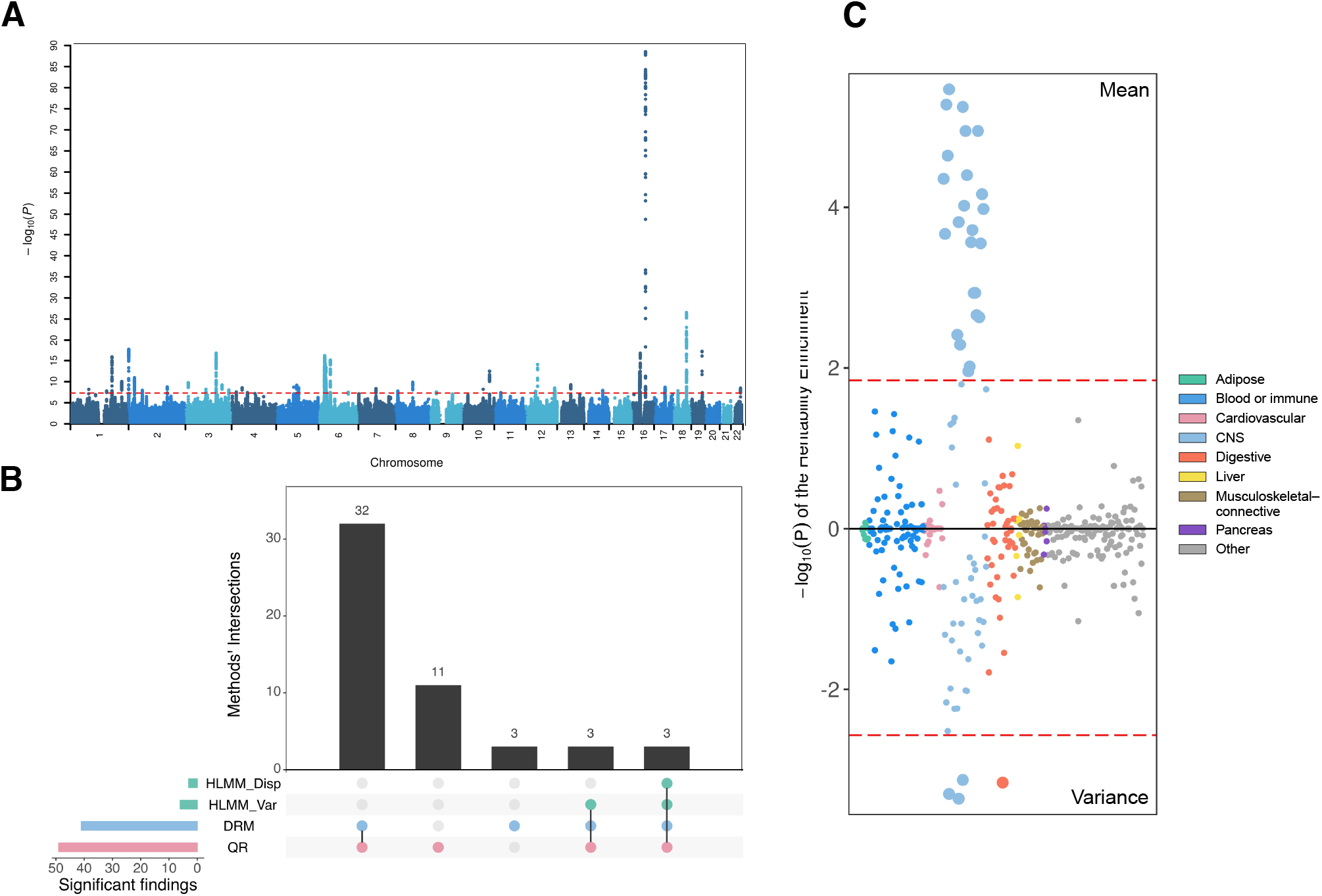
vQTL mapping of BMI in UK Biobank. **(A)** Manhattan plot of genome-wide vQTL analysis for BMI in UK Biobank using QUAIL. The dashed red line indicates P = 5.0e-8. **(B)** Number of independent significant loci (P < 5.0e-8) identified by four vQTL methods. This plot uses bars to break down the Venn diagram of overlapped loci in different vQTL methods. **(C)** Cell-type enrichment results for BMI vQTL (upper) and GWAS associations (lower). Each data point represents a tissue or cell type. Different colors represent tissue categories based on Finucane et al^31^. Dashed red lines are drawn at FDR = 0.05.

Previous studies have shown that heritability of BMI is mostly enriched in active genomic regions of the central nervous system (CNS)^31,32^. A recent study showed that vQTL of BMI are significantly enriched in the gastrointestinal tract^13^. We applied stratified LD score regression^33^ to summary statistics of QUAIL vQTL and GWAS of BMI. We partitioned the vQTL and GWAS associations by 205 cell-type-specific annotations^34,35^. Overall, we observed similar cell type enrichment patterns between GWAS and vQTL associations (Pearson’s correlation of LD score regression coefficient across 205 annotations = 0.78, P = 2.1e-71; **Supplementary Table 2**). Both vQTL and GWAS signals showed strong enrichment in CNS. The stomach cell type was specifically enriched for BMI vQTL (**Figure 3C**, P = 6.9e-4) but not GWAS heritability (P = 0.22), suggesting different biological mechanisms underlying the level and variability of BMI.

### GxE enrichment in vQTL

To investigate whether BMI vQTL are enriched for GxE interactions, we performed GxE interaction tests using genome-wide SNP data and two BMI-related behavioral traits in UK Biobank: physical activity (PA)^3,36,37^ and sedentary behavior (SB)^15,38^. We assessed enrichment for nominally significant GxE interactions (P < 0.05) in top vQTL and GWAS associations for BMI (**Supplementary Table 3**; **Methods**). We observed consistently and substantially stronger enrichment for GxPA and GxSB interactions in top vQTL than in top GWAS associations for BMI (**Figure 4**). These results show that vQTL mapping may be a more effective strategy to screen for GxE candidates than GWAS. In addition, although the fold enrichment has a decreasing trend as we consider more vQTL in the analysis, we still observed substantial and highly significant GxE enrichment even in top 15% of vQTL for both PA (fold enrichment = 1.66, P = 4.0e-109) and SB (fold enrichment = 1.51, P = 1.5e-87), suggesting pervasive GxE interactions among SNPs associated with BMI variability.

**Figure 4.**
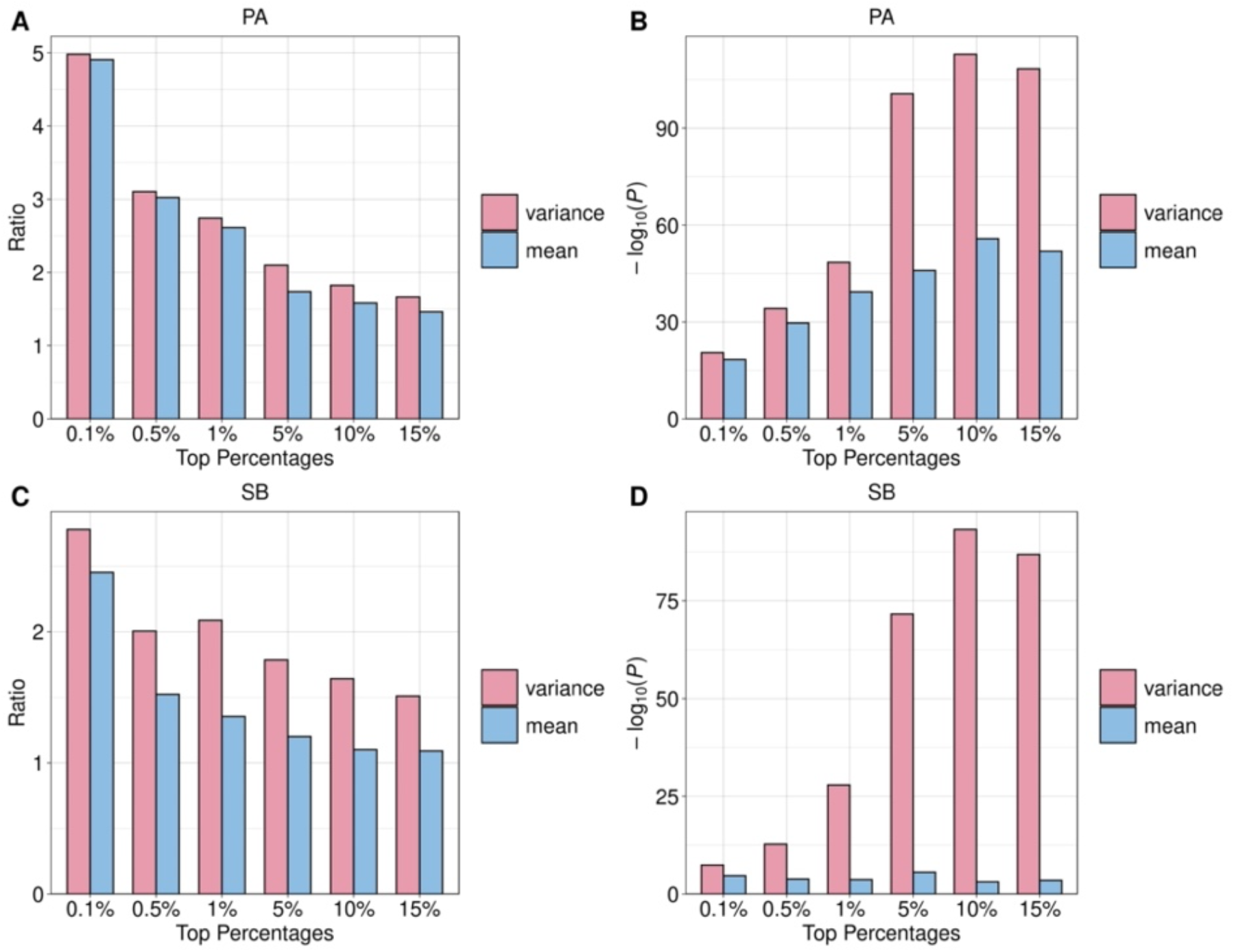
Enrichment for GxE interactions in top BMI vQTL and GWAS associations. Panels **A** and **C** illustrate the fold enrichment for GxE interactions in top vQTL (pink) and GWAS associations (blue). Fold enrichment ratio is defined as the actual count of significant GxE among top SNPs divided by the expected count. Panels **B** and **D** illustrate the p-values for enrichment calculated from Fisher’s exact test. The environmental factors are physical activity (PA) for Panels **A-B**, and sedentary behavior (SB) for **C-D**.

### vPGS predicts population-level and within-individual variability of BMI

Next, we explore if genome-wide vQTL associations can be aggregated into concise, effective metrics to better quantify genetic effects on trait variability. Although it is straightforward to generate vPGS using vQTL effect sizes as SNP weights, it is a non-trivial task to evaluate the predictive performance of vPGS. Common metrics that are used to assess PGS performance (e.g., R^2^) quantify association between PGS and trait levels and do not reflect the effect of vPGS on trait variability. Here, we extend our quantile regression framework to continuous predictors (**Methods**) and use it to benchmark the performance of different vPGS models.

We first investigated if vPGS can predict the population-level BMI variability (**Supplementary Figure 3**) using three independent longitudinal datasets, i.e., Health and Retirement Study (HRS), Wisconsin Longitudinal Study (WLS), and National Longitudinal Study of Adolescent to Adult Health (Add Health). We describe details of sample QC procedures in **Methods**. We used a multi-level linear growth curve model to adjust for age effects on longitudinal measurements of BMI. In each longitudinal cohort, we estimated the expected BMI of each individual across waves after removing age effects (**Methods**). We generated vPGS in each cohort using vQTL effects estimated in UK Biobank by QUAIL, HLMM_Var, HLMM_Disp, and DRM. vPGS based on QUAIL vQTL consistently showed the largest effect sizes and the most significant associations with BMI variability in three independent cohorts (**Table 2**), followed by DRM. Compared with individuals in the lowest vPGS quintile, individuals in the highest quintile showed 61%, 52%, and 73% increase in BMI variance in HRS, Add Health, and WLS, respectively (**Figure 5A**). We also obtained similar results using double generalized linear model (DGLM) as an alternative approach to evaluate vPGS performance (**Methods**; **Supplementary Table 4**), with vPGS based on QUAIL consistently showing the strongest associations with BMI variability.

**Table 2.** Benchmarking the prediction accuracy of vPGS for population-level and within-individual BMI variability. The upper and lower tables show the results of population-level and within-individual variability, respectively. Each row represents a different vPGS approach. In the upper table, Beta denotes the estimated effect size of vPGS on the population-level BMI variability using an evaluation method based on our quantile regression approach. In the lower table, Beta denotes the estimated effect of vPGS on the coefficient of variation (CV). SE is the standard error of estimated effects. The most predictive vPGS is highlighted in boldface.

| Population-level variability |  |  |  |  |  |  |  |  |  |
| --- | --- | --- | --- | --- | --- | --- | --- | --- | --- |
| Methods | HRS(N=10,550) |  |  | Add Health (N=6,717) |  |  | WLS (N=4,694) |  |  |
|  | Beta | SE | P-value | Beta | SE | P-value | Beta | SE | P-value |
| QUAIL | **0.520** | **0.055** | **3.07E-21** | **0.716** | **0.090** | **1.89E-15** | **0.610** | **0.077** | **2.16E-15** |
| HLMM_Var | 0.260 | 0.058 | 7.08E-6 | 0.268 | 0.094 | 0.106 | 0.292 | 0.084 | 4.93E-4 |
| HLMM_Dis | 0.098 | 0.058 | 0.094 | -0.020 | 0.094 | 0.833 | 0.156 | 0.086 | 0.068 |
| DRM | 0.507 | 0.059 | 9.97E-18 | 0.644 | 0.093 | 3.66E-12 | 0.521 | 0.082 | 2.33E-10 |
| Within-individual variability |  |  |  |  |  |  |  |  |  |
| Methods | HRS (N= 10,502) |  |  | Add Health (N= 6,706) |  |  | WLS (N= 4,471) |  |  |
|  | Beta | SE | P-value | Beta | SE | P-value | Beta | SE | P-value |
| QUAIL | **0.097** | **0.010** | **9.31E-24** | **0.092** | **0.012** | **2.60E-14** | **0.088** | **0.015** | **2.28E-9** |
| HLMM_Var | 0.048 | 0.010 | 4.26E-7 | 0.048 | 0.012 | 8.70E-5 | 0.048 | 0.015 | 0.012 |
| HLMM_Dis | 0.020 | 0.010 | 0.035 | 0.014 | 0.012 | 0.236 | 0.034 | 0.015 | 0.021 |
| DRM | 0.086 | 0.010 | 6.47E-17 | 0.087 | 0.012 | 2.07E-12 | 0.082 | 0.015 | 8.47E-8 |

**Figure 5.**
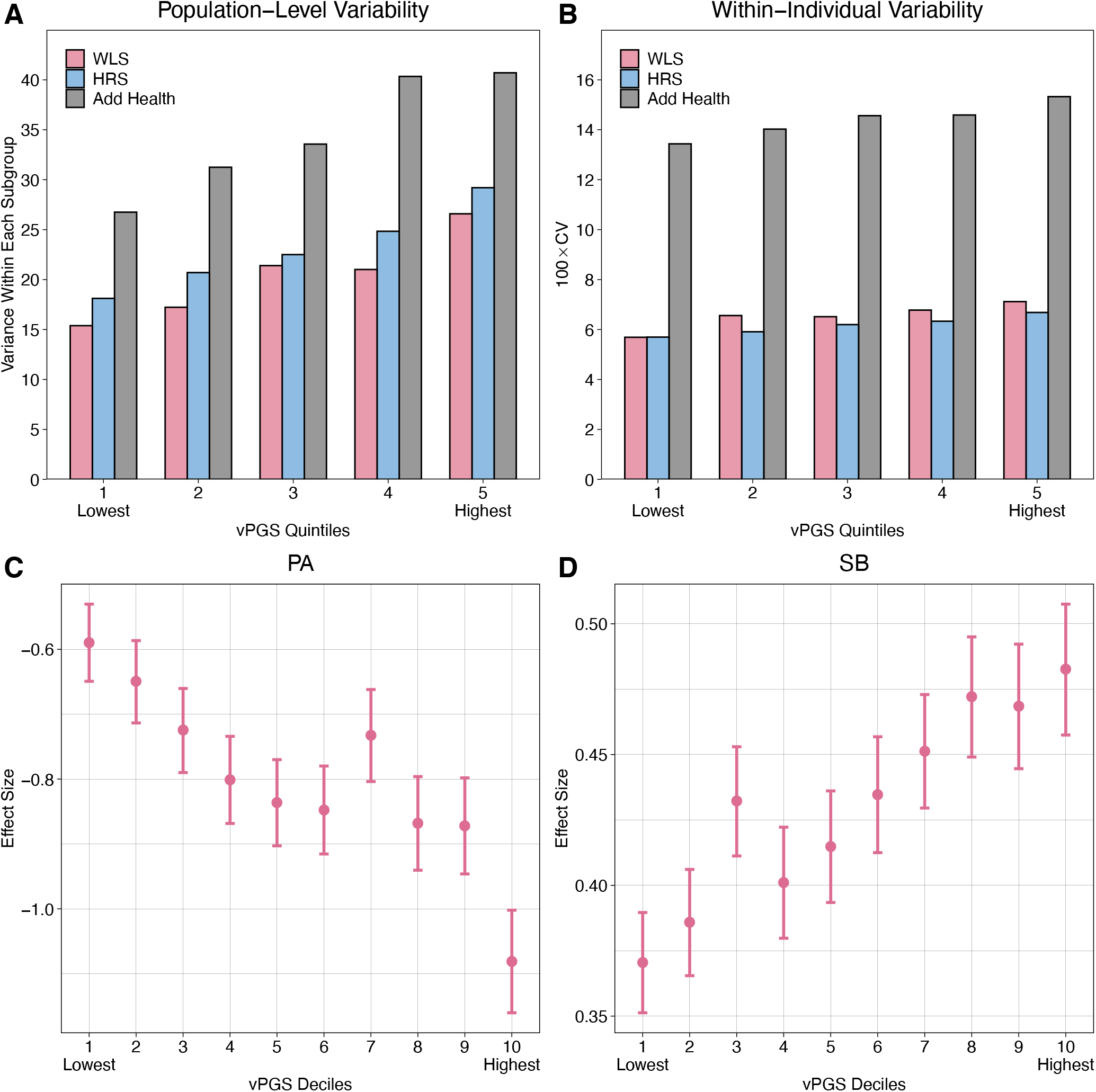
vPGS performance and application in GxE interaction. Panels **A-B** illustrate the prediction accuracy of vPGS on population-level and within-individual BMI variability, respectively, in three external cohorts. **(A)** Each bar shows the variance of BMI within each vPGS quintile in a given cohort. **(B)** Each bar shows the average within-individual BMI variability quantified by the 100×Coefficient of Variation (CV) within each vPGS quintile. Panels **C-D** illustrate the vPGS-PA and vPGS-SB interactions in UK Biobank holdout samples. **(C)** The effect size of PA on BMI by vPGS deciles. **(D)** The effect size of SB on BMI by vPGS deciles.

We continued to investigate if vPGS estimated from cross-sectional data is also predictive of within-individual variability which quantifies the change in a dynamic outcome (e.g., BMI) as individuals progress through the life course (**Supplementary Figure 3**). Although within-individual trait variability is a better way to quantify outcome plasticity in response to environmental changes, direct estimation of genetic associations with within-individual variability remains challenging, mostly due to limited samples in existing cohorts with genotype data and longitudinal phenotypic measurements. We leveraged the longitudinal nature of the three datasets described above and used the wave-to-wave variability to quantify the within-individual variability. More specifically, we estimated the wave-to-wave BMI variability using the coefficient of variation (CV; **Methods**). To benchmark the performance of vPGS, we used linear regressions to quantify vPGS associations with CV in each cohort. vPGS based on QUAIL again showed the best predictive performance among all methods, followed by DRM (**Table 2**). vPGS based on HLMM showed substantially weaker associations with CV in all cohorts. **Figure 5B** shows the average within-individual CV for samples in each vPGS quintile. We observed 17%, 14%, and 25% increase in within-individual BMI variability in the highest vPGS quintile than in the lowest quintile for HRS, Add Health, and WLS, respectively.

### GxE interaction analysis using vPGS

To further investigate the possibility of using vPGS in GxE interaction studies, we randomly apportioned unrelated UKB participants of European descent into training and testing sets with an 80-20 split. We first applied QUAIL to estimate vQTL effects of all SNPs on BMI using samples in the training set, and then used QUAIL summary statistics to generate vPGS for samples in the testing set. We tested vPGS-PA and vPGS-SB interactions for BMI in the testing samples (**Methods**).

We identified significant interactions between BMI vPGS and both PA (P = 1.1e-8) and SB (P = 1.6e-5) (**Supplementary Table 5**). Both interactions remained significant after adjusting for vPGS-covariate interaction terms in the model^39^ (P = 1.7e-8 and 1.1e-7 for PA and SB, respectively; **Supplementary Table 6**). We partitioned the testing sample into 10 deciles based on vPGS values and observed clear, linearly decreasing trajectories of PA effects and increasing SB effects on BMI as vPGS increases (**Figures 5C** and **5D**).

## Discussion

In this paper, we introduced QUAIL, a novel, unified statistical framework for estimating genetic effects on the variability of quantitative traits. QUAIL constructs a quantile integral phenotype which aggregates information from all quantile levels, and only requires fitting two linear regressions per SNP in genome-wide analysis. Our approach directly addresses some limitations of current vQTL methods, including a lack of robustness to non-Gaussian phenotypes and confounding effects on both trait levels and trait variability. We also demonstrated that QUAIL can be extended to continuous predictors such as vPGS. Applied to 375,791 samples in UK Biobank, QUAIL identified 49 significant vQTL for BMI, including 11 novel loci that have not been previously identified. These vQTL were significantly enriched in functional genomic regions in CNS and gastrointestinal tract, were substantially enriched for GxE interactions with BMI-related behavioral traits, and produce vPGS that can effectively predict both population-level and within-individual BMI variability. Overall, these results hinted at distinct genetic mechanisms underlying the level and variability of BMI.

Evidence suggests that genetics, environments, and their ubiquitous interactions jointly shape human phenotypes^1^. However, there has only been limited success in identifying robust GxE interactions in complex trait research. This is because detecting GxE interactions at the SNP level requires a hypothesis-free genome-wide scan which introduces an extreme burden of multiple testing and severely reduces statistical power. Alternatively, people constructed PGS which are genome-wide summaries of numerous SNPs’ aggregated effects on trait levels and used these scores as the G component in GxE studies^5-8^. However, these scores do not directly quantify the susceptibility of each individual to environmental exposures and could only partially characterize the interplay between genes and environments. Our study advances the field on multiple fronts. First, our approach produces statistically robust and powerful vQTL results. These loci associated with phenotypic variability may be used as candidate SNPs in GxE research, thereby reducing the search space for possible interactions. Second, we demonstrated that vPGS based on QUAIL effect estimates show superior predictive performance compared to existing approaches. The improved vQTL and vPGS, coupled with large population cohorts with deep phenotyping and sophisticated measurements on the environments, have the potential to improve prioritization and aggregation of genetic effects on both trait levels and plasticity and accelerate findings in GxE research.

Our study has some limitations. First, our method cannot be applied to binary phenotypes. Second, it is unclear if a linear mixed model accounting for sample relatedness will be compatible with the quantile integral phenotype produced by QUAIL. Third, the use of vPGS to predict the within-individual phenotypic variability requires some attention. In the paper, we generated vPGS using the vQTL effects obtained from a genome-wide analysis of population-level BMI variability and demonstrated its significant association with the longitudinal, wave-to-wave BMI variability. However, for certain traits, it is possible that within-individual and population-level variability are controlled by distinct biological processes and have different genetic architecture. An ultimate solution to studying the genetic basis of within-individual variability requires large GWAS samples with repeated measurements of the same outcome for each individual across time. Finally, it is known that genetic effects on the level and the variability of BMI can be correlated^12,13,16^. We also made similar observations in our analysis (**Supplementary Figure 4**). Young et al.^16^ previously introduced dispersion effect which quantifies the residual genetic effect on trait variance after de-correlating association with trait levels. But this approach may be overly conservative especially when SNP-trait associations are heteroskedastic. It also requires an inverse-normal transformation to the phenotype which has been suggested to reduce GxE signals^13^. In the **Supplementary Note**, we show that dispersion effect can also be estimated in our framework. It substantially reduces the mean-variance relationship (**Supplementary Figure 4**) but identifies fewer loci for BMI (**Supplementary Figures 5-6** and **Supplementary Table 7**). When and how to use these dispersion effect estimates in GxE applications remains to be explored in the future.

Taken together, QUAIL addresses several critical limitations in existing vQTL and vPGS methods and provides robust, powerful, and computationally efficient estimates for genetic effects on phenotypic variability. These methodological advances, in conjunction with increasing sample size in population cohorts with longitudinal measures of phenotypic outcomes and environments, promise exciting new developments in the near future. We believe our approach complements existing analytical strategies and will have broad applications in future studies of complex trait genetics and GxE interactions.

## Methods

### Statistical model

If a SNP *G* is a vQTL for trait *Y*, the slopes (i.e., *β* _*τ*_) will differ in quantile regressions based on different quantile levels *τ* (**Figure 1**).

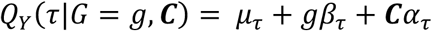

Here, ***C*** is a *n* × *m* matrix for covariates in the model, *α*_*τ*_ denotes the regression coefficients for covariates, and *μ*_*τ*_ is the intercept. For a pair of quantile levels (1 − *τ, τ*), *τ*∈ (0, 0.5), the difference between regression coefficients (i.e., *β*_1−*τ*_ − *β* _*τ*_) quantifies the effect of SNP on the variability of *Y*. Instead of choosing arbitrary quantile levels to define the effect size, we aggregate information across all quantile levels to define the vQTL effect:

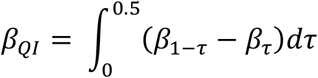

Note that more generally we can use ∫*w*_*τ*_*βτdτ* to quantify the effect. In this study, we set *w*_*τ*_ = 1 when *τ*≥ 0.5 and *w*_*τ*_ = −1 when *τ*< 0.5. Testing if a SNP is associated with the variability of *Y* is equivalent to testing the null *H*_0_: *β*_*QI*_ = 0. In practice, *β*_*QI*_ can be approximated using a linear spline expansion from *K* quantile levels:

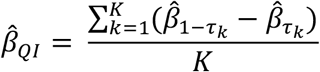

There are two key inference problems in this framework. First, to obtain parameter estimates 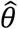 which include 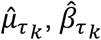, and 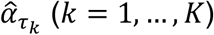 (*K* = 1, …, *K*), we can use a standard fitting approach for quantile regression^40^:

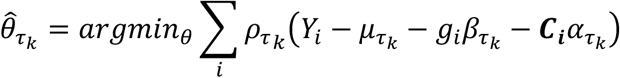

where 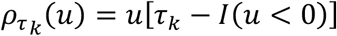 and *i* is the index for the *i* -th individual in the analysis. However, to make the linear spline approximation accurate for *β*_*QI*_, *K* needs to be big. This will lead to fitting *K* quantile regressions for each SNP which is computationally challenging in genome-wide analysis. Second, the standard error for quantile integrated effect 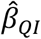 involves estimation of the variance-covariance matrix for 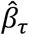 and is difficult to obtain. We propose a two-step procedure in QUAIL to obtain statistically justified estimates for quantile integral effect while bypassing these computational challenges.

**Step 1:** Transform the phenotype into a quantile integrated rank score.

First, we estimate the intercept 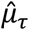 and covariate effects 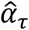 under the null model (i.e., *β* _*τ*_ = 0) for 2*K* quantile levels

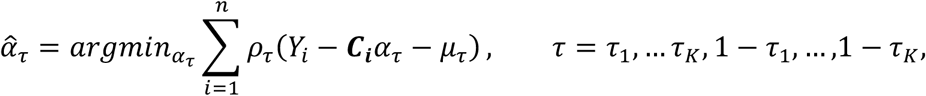

where *ρ*_*τ*_ (*u*) = *u*[*τ*− *I*(*u* < 0)] is the loss function for quantile regression. Importantly, this step is done on the null model, so it does not need to be repeated for different SNPs in genome-wide analysis. Then, for each individual *i*, we construct 2*K* quantile rank scores:

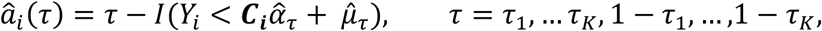

where 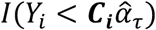 is a binary indicator for whether *Y*_*i*_ is smaller than the estimated *τ*^*th*^ conditional quantile for *Y*_*i*_.

Then, we construct a quantile rank score for each individual:

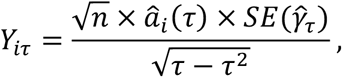

where *n* is the sample size, 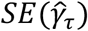 is the standard error of the regression coefficient estimate 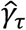 in quantile regression *QY*(*τ*|*d*, ***C***) = *b*_*τ*_ + *dγ*_*τ*_ + ***C****α*_*τ*_ and *d* is a random variable sampled from *N*(0,1). Here, we create the random variable *d* and calculate 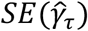 as described above to approximate 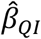 using estimated quantile regression coefficients 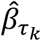 while bypassing the fitting of *K* quantile regressions for each SNP. We show the details and rationale of this approximation in **Supplementary Note** and **Supplementary Figure 7-8**.

Finally, we construct the quantile integrated rank score for each individual *i* by combining *Y*_*iτ*_ across quantile levels:

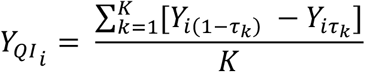

We then center the 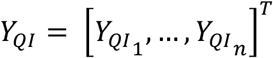 to have a mean 0.

**Step 2:** Estimate the quantile integral effect.

We estimate the quantile integral effect as

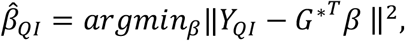

where *G*^*^ is the *n* × 1 vector of genotype residuals after regressing out covariates. More specifically, *G*^*^ = (*I* -*P*_*C*_)*G*, where *G* is the original *n* × 1 standardized genotype vector with mean 0 and variance 1, *C* is the *n* × *m* matrix for covariates, and *P*_*C*_ =*C*(*C*^*T*^*C*) ^−1^*C*^*T*^ is the projection onto the linear space spanned by *C*. Since we adjusted for covariates when obtaining the *Y*_*QI*_ and *G*^*^, the quantile integral effect is account for the covariates’ effects on traits level and variance. We provide detailed derivations of this procedure in the **Supplementary Note**.

Under the null hypothesis that the slopes (i.e., *β* _*τ*_) are identical in quantile regressions based on different quantile levels 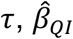 follows a normal asymptotic distribution

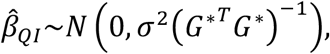

where *σ*^2^ = *Var*(*ϵ*) and *ϵ* is the residual in linear regression *Y*_*QI*_ = *G*^*^*β* + *ϵ*. We provide the derivation for the null distribution of test statistics in the **Supplementary Note**. In our implementation, we use a linear regression *Y*_*QI*_ = *G*^*^*β* + *ϵ* to obtain the QUAIL test statistics and p-values.

### Simulation settings

We performed extensive simulations to evaluate the type-I error, statistical power, and the ability to correct for confounding effect on trait variability for five vQTL methods including QUAIL, LT, DRM, and HLMM with and without inverse normal transformation. We used 100 quantile levels (i.e., *K* = 100) for QUAIL. We generated a SNP variable *G* coded as 0, 1, 2 from *Binomial*(2, *f*), where *f* is the minor allele frequency (MAF) generated from a uniform distribution on [0.05, 0.5]. Environmental exposure *E* was generated from a standard normal distribution *N*(0,1). We repeated the simulation 1000 times and calculated FPR and power as the proportion of simulations where the null hypothesis was rejected at *P* < 0.05.

For FPR simulations, we used a model where the SNP only has effects on the level but not the variance of the phenotype. We simulated phenotype for 10,000 individuals according to the model *y*_*i*_ = *β*_*g*_*G*_*i*_ + *ϵ*_*i*_, *i* = 1, …,10000, where *y*_*i*_ is the simulated phenotype, *ϵ*_*i*_ is an error term with mean 0 and variance 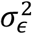 for the *i*-th individual. To simulate the error term with different levels of skewness and kurtosis, we sampled ϵ_*i*_ from three different distributions: standard normal distribution *N*(0, 1), t distribution with df = 3, and *χ*^2^ distribution with df = 6. Regression coefficients were selected such that the proportion of total PVE by genotype, defined as *Var*(*β*_*g*_*G*_*i*_)/*Var*(*y*_*i*_) ranged between 0.5% to 5%. 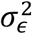 was set to be 1 − *Var*(*β*_*g*_ *G*_*i*_)/*Var*(*y*_*i*_) so that *Var*(*y*_*i*_) = 1.

For power simulation, we simulated the phenotype such that the SNP only has a variance effect on the phenotype. This variance effect is reflected in the interaction term for the SNP and environmental exposures. We simulated phenotype for 10,000 individuals according to the model *y*_*i*_ = *β*_*GE*_*G*_*i*_*E*_*i*_ + ϵ_*i*_, *i* = 1, …,10000, where *y*_*i*_ is the simulated phenotype, ϵ_*i*_ is an error term with mean 0 and variance 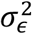 for the *i* -th individual. We also simulated error terms from three different distributions as described above. We selected the regression coefficients such that the proportion of total PVE by the GxE interaction, i.e., *Var*(*β*_*GE*_*G*_*i*_*E*_*i*_)/*Var*(*y*_*i*_), ranged between 0.5% to 5%. We set 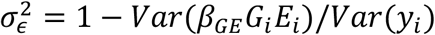.

To assess different methods’ robustness to confounding effect on trait variability, we simulated the phenotype such that the SNP has no effect and a covariate has variance effect on the phenotype. This covariate was generated from *Bernoulli*(*p*) where *p* varies with each individual’s genotype value (i.e., *p* = 0.2 for individuals with *G* = 0, *p* = 0.5 when *G* = 1, and *p* = 0.8 when *G* = 2). The covariate’s variance effect is reflected in the interaction term for the covariate and environmental exposures. We simulated phenotype for 10,000 individuals according to the model *y*_*i*_ = *β*_*CE*_*C*_*i*_*E*_*i*_ +*ϵ*_*i*_, *i* = 1, …,10000, where *y*_*i*_ is the simulated phenotype, *C*_*i*_ is the covariate, ϵ_*i*_ is an error term that follows 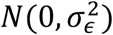 for the *i* -th individual. We selected the regression coefficients such that the proportion of total PVE by the covariate x E interaction, i.e.,*Var*(*β*_*CE*_*C*_*i*_*E*_*i*_)/*Var*(*y*_*i*_), ranged between 2.5% to 20%. We also set 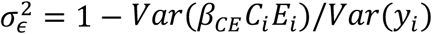 to rescale the variance of *y*_*i*_ to be 1.

### UK Biobank data processing

QC procedure of genetic data in the UK Biobank has been described elsewhere^41^. We analyzed UK Biobank samples with European ancestry inferred from genetic PCs (data field 22006). Participants that are recommended by UK Biobank to be excluded (data field 22010), those with conflicting genetically inferred (data field 22001) and self-reported sex (data field 31), and those who withdrew from the study were excluded from the analyses. We also removed related individuals identified by KING^42^ and retained 377,509 unrelated individuals with European descent.

### Genome-wide vQTL mapping for BMI

Following previous work on genome-wide analysis in UK Biobank^43^, we used year of birth (data field 34), sex (data field 31), genotyping array, and top 12 PCs computed using flashPCA2^44^ on the analytical sample as covariates for both trait level and variability. We only included SNPs with a MAF > 1% and missingness < 1% in the analysis.

We conducted genome-wide vQTL analysis using four methods: QUAIL, DRM, and HLMM_Var, and HLMM_Disp. For QUAIL, we transformed the BMI into a quantile integrated rank score, obtained SNP residual values by regressing each SNP on covariates, and estimated the vQTL effect by regressing the quantile integrated rank score on SNP residuals. For DRM, we first fit a linear model between BMI and covariates and calculated the BMI residual. Then, we applied DRM to quantify the effect of each SNP on BMI residual. For HLMM, we first applied an inverse normal transformation to BMI. Then, we fit HLMM to obtain the additive and log-linear variance effects (i.e., HLMM_Var). Next, we estimated the HLMM dispersion effect (i.e., HLMM_Disp) by using the additive and log-linear variance effects as described previously^16^.

We set the genome-wide significance threshold as 5.0e-8. To determine the number of independent significant vQTL, we clumped the summary statistics for each method in PLINK2^45^ (--clump option with parameters --clump-p1 5.0e-8 --clump-p2 5.0e-8 --clump-r2 0.01 and --clump-kb 5000) using the analytic sample in UK Biobank as the LD reference panel. To visualize the results, we generated the Manhattan plot and quantile-quantile plot using the ramwas^46^ package in R.

We also conducted a GWAS for BMI using Hail^47^ on the same data used in the vQTL analysis. We included year of birth, sex, genotyping array, and top 12 PCs computed using flashPCA2^44^ on the analytical sample as covariates. LD clumping and visualization were performed similarly as described above.

Additionally, we used the estimated intercept from LD score regression^48^ to quantify the level of unadjusted confounding in genome-wide vQTL analysis. We used ashR^30^ on the full set of SNPs to estimate the proportion of non-null vQTL associations.

### Cell-type heritability enrichment analysis

We used stratified LD score regression^33^ to perform cell-type enrichment analyses with gene expression data using the “Multi_tissue_gene_expr” (including data from GTEx and Franke lab) flag and default settings. We only included non-MHC HapMap3 SNPs for LD score regression analysis. Cell-type enrichment p-values across 205 functional annotations were adjusted using the Benjamini-Hochberg method for false discovery rate^49^.

### Gene-environment interaction enrichment analysis

We performed GxE interaction tests using genome-wide SNP data and two BMI-related behavioral traits (i.e., PA and SB) in UK Biobank. Details about the construction of PA and SB variables can be found elsewhere^13,15^. For PA, we assigned a three-level categorical score (low, medium, and high) based on the short form of the International Physical Activity Questionnaire (IPAQ) guideline for each individual. We defined SB as an integer using the combined time (hours) spent driving, using computer, and watching television.

To assess the enrichment for GxE effects in top vQTL, we first clumped the QUAIL summary statistics in PLINK2 (--clump option with parameters --clump-p1 1 --clump-p2 1 --clump-r2 0.1 and --clump-kb 1000) using the CEU samples in 1000 Genome Project Phase III cohort as the LD reference panel. Next, we performed a GxE analysis to test the interaction between each SNP in the clumped summary statistics and PA and SB based on the model:

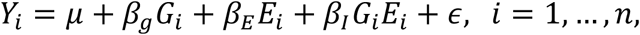

where *Y*_*i*_ is BMI, *G*_*i*_ is the SNP genotype, and *E*_*i*_ is the environmental factor for the i-th individual. We defined nominally significant GxE using a p-value cutoff of 0.05. We also defined vQTL as the top 0.1, 0.5, 1, 5, 10, and 15 percent of SNPs ordered by their QUAIL p-values in the clumped summary statistics. The fold enrichment is calculated as the actual count of significant GxE among vQTL divided by the expected count. We used Fisher’s exact test to test the enrichment.

For comparison, we also performed enrichment analysis for GxE interaction in top GWAS associations using the same analytical procedure described above.

### Predicting population-level trait variability

To benchmark the predictive power of vPGS, we used data from three independent cohorts: HRS, Add Health, and WLS. We only included individuals of European ancestry in the analysis. The sample size is 10,550, 6,717, and 4,694 for HRS, Add Health, and WLS respectively.

To compute vPGS, we first clumped each set of summary statistics in PLINK2 (--clump option with parameters --clump-p1 1 --clump-p2 1 --clump-r2 0.1 and --clump-kb 1000) using the CEU samples in 1000 Genome Project Phase III cohort as the LD reference panel. Then, we computed vPGS using PRSice-2^50^ without p-value filtering. We calculated four vPGS based on different vQTL methods: QUAIL, DRM, HLMM_Var, and HLMM_Disp.

To quantify the performance of vPGS in predicting the population-level variability, we first fit a multi-level linear growth curve model on BMI and age in each cohort:

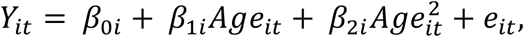

where *Y*_*it*_ and *Age*_*it*_ denote the BMI and age of respondent *i* at time point *t*, respectively (*i* = 1, …, *n* and *t* = 1, …, *T*_*i*_), *β*_0*i*_ is assumed to be normally distributed. We included linear and quadratic terms for age to reflect the non-linear age-dependent trajectory of BMI. The estimated individual intercept (i.e.,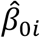) represents the expected BMI after removing age effect. We denote it as BMI-adj and use it as the trait value for the further analysis described below.

We extended our quantile regression framework to assess vPGS performance with two modifications due to the eased computational burden. First, we regress the phenotype on the vPGS and use the residual to construct the rank score *â*_*i*_(*τ*) as described before. Second, we perform a standard quantile regression and use the kernel-based sandwich approach^51^ to obtain the standard error of the estimated quantile regression coefficient for vPGS, i.e.,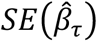. We use the same approach to construct the quantile integrated rank score *Y*_*QI*_ for each individual. The effect size of vPGS can be quantified as

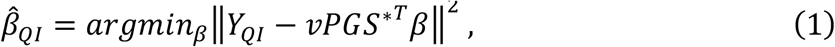

where *vPGS*^*^ is the *n* dimensional vPGS residual vector after regressing out covariates. Here, original vPGS is standardized to have mean 0 and variance 1. We use this quantile integral effect to quantify the predictive performance of vPGS on the population-level variability. We adjusted sex and top 10 PCs in the analysis of each cohort.

We also extend the DGLM^14,26^, the method implemented in HLMM, as an alternative approach to evaluate vPGS performance. The DGLM takes the form of

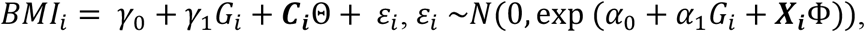

where *BMI*_*i*_ denotes the inverse normal-transformed BMI-adj of individual *i, G*_*i*_ is the vPGS of individual *i, **C***_*i*_ is the vector of covariates including sex and top 10 PCs. Here, *α*_1_ quantifies the effect of vPGS on the variability of BMI and is the parameter of interest in this analysis. We fitted DGLM using the dglm^52^ packages in R.

To visualize the predictive performance of vPGS in predicting the population-level variability, we divided samples into 5 quintiles according to their vPGS values and compared the variance of BMI-adj across quintiles in each cohort.

### Predicting within-individual trait variability

We used the same three external datasets (i.e., WLS, HRS, and Add Health) to benchmark the performance in predicting within-individual BMI variability. We applied the same QC procedure described above except that we only included individuals with reported BMI in at least two waves.

We quantified the wave-to-wave BMI variability using CV^53^ defined as:

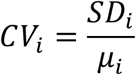

where *SD*_*i*_ is the BMI standard deviation of the *i*-th individual across waves and *μ*_*i*_ is the individual mean of BMI across waves. We calculated CV for each individual based on all of the participant’s BMI measurements across waves. Then, we used linear regression to quantify the predictive performance of vPGS on the within-individual variability. We regressed CV on vPGS in each cohort and included sex, mean age across waves, and top 10 PCs as covariates. To visualize the results, we divided samples from each cohort into 5 quintiles according to their vPGS values and compared the average CV across quintiles.

### Gene-environment interaction analysis using vPGS

To test GxE interactions using vPGS, we randomly apportioned unrelated UKB participants of European descent (N=375,791) into training (N=300,633) and testing sets (N=75,158), with an 80-20 split. We applied QUAIL to estimate the effect of each SNP on BMI variability using samples in the training set, while controlling for year of birth, sex, genotyping array, and top 12 PCs computed using flashPCA2^44^ on the analytical sample as covariates.

Next, we used weights obtained in the training set to construct vPGS for samples in the testing set. To compute vPGS, we first clumped the summary statistics in PLINK2 (--clump option with parameters --clump-p1 1 --clump-p2 1 --clump-r2 0.1 and --clump-kb 1000) using the CEU samples in 1000 Genome Project Phase III cohort as the LD reference panel. Then, we computed vPGS using PRSice-2 without p-value filtering.

We tested vPGSxE effects on BMI by fitting the following model:

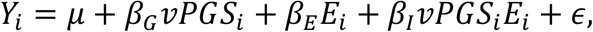

where *Y*_*i*_ is BMI, *vPGS*_*i*_ is the standardized vPGS with mean 0 and variance 1, and *E*_*i*_ is the environmental factor (i.e., PA or SB) for the *i*-th individual. We adjusted for year of birth, sex, genotyping array, and top 12 PCs. To check the robustness of our results, we repeated our vPGSxE analysis on BMI using the model above with vPGS-Sex and vPGS-Year of birth interaction terms as additional covariates.

To visualize the interaction, we divided samples into 10 deciles based on their vPGS values and compared estimates of the environmental factor on BMI across vPGS deciles.

### URL

UK Biobank (http://www.ukbiobank.ac.uk/);

HRS (https://hrs.isr.umich.edu/about);

Add Health (https://addhealth.cpc.unc.edu/);

WLS(https://www.ssc.wisc.edu/wlsresearch/);

HLMM (https://hlmm.readthedocs.io/en/latest/);

DRM (https://github.com/drewmard/DRM); ashR (https://github.com/stephens999/ashr).

## Data and code availability

QUAIL software package is publicly available at (https://github.com/qlu-lab/QUAIL). Summary statistics of QUAIL vQTL analysis for BMI are available at (http://qlu-lab.org/data.html).

## Acknowledgments

This project was supported by the pilot grant of the Center for Demography of Health and Aging at University of Wisconsin-Madison (P30 AG017266). We also acknowledge research support from the University of Wisconsin-Madison Office of the Chancellor and the Vice Chancellor for Research and Graduate Education with funding from the Wisconsin Alumni Research Foundation. We thank faculties and students involved in the Initiative in Social Genomics at the University of Wisconsin-Madison for helpful discussions.

## Author contribution

Q.L. conceived and designed the study.

J.M. and Q.L. developed the statistical framework.

J.M. performed simulations and data analysis.

Y.L. assisted in literature review and model conceptualization.

Y.W. assisted UK Biobank data preparation.

B.Z and J.F. assisted HRS and Add Health data preparation.

L.S. advised on GxE interaction.

Q.L. advised on statistical and genetic issues.

J.M. and Q.L. wrote the manuscript.

All authors contributed in manuscript editing and approved the manuscript.

