## Supplementary Note and Figure for "A quantile integral linear model to quantify genetic effects on phenotypic variability"

### Rationale for using quantile regression to prioritize GxE loci

Consider a linear model with main effects for genetics and environments and their interaction

$$Y_i = \beta_0 + \beta_G G_i + \beta_E E_i + \beta_I G_i E_i + \epsilon_i\,, \quad i = 1, \dots, n,$$

where $Y_i$ is the quantitative phenotype, $E_i$ is the environmental factor following a distribution $F_E$ with mean $\mu_E$ and variance $\sigma_E^2$ (i.e., $E_i \sim F_E(\mu_E, \sigma_E^2)$), $G_i$ is the additive SNP genotype with values 0, 1, or 2, and $\epsilon_i$ is the error term following a distribution $F_\epsilon(0, \sigma_\epsilon^2)$. We assume $G_i$, $E_i$, and $\epsilon_i$ to be mutually independent. Under this model, the conditional expectation and variance of $Y_i$ given the genotype $G_i = g_i$ are

$$E[Y_i|G_i = g_i] = (\beta_0 + \beta_E\mu_E) + (\beta_G + \beta_I\mu_E)g_i,$$
$$Var[Y_i|G_i = g_i] = \sigma_\epsilon^2 + (\beta_E + \beta_I g_i)^2\sigma_E^2,$$

and we also have the following conditional heteroscedastic linear model

$$Y_i|g_i = (\beta_0 + \beta_E\mu_x) + (\beta_G + \beta_I\mu_x)g_i + (\beta_E + \beta_I g_i)\epsilon_{1i} + \epsilon_{2i}\,,$$

where $\epsilon_1 \sim F_{\epsilon_1}(0, \sigma_E^2)$ and $\epsilon_2 \sim F_{\epsilon_2}(0, \sigma_\epsilon^2)$.

Then, we can derive the conditional quantile regression function

$$Q_Y(\tau|G_i = g_i) = \left[\beta_0 + \beta_E\mu_x + \beta_E Q_{\epsilon_1}(\tau) + Q_{\epsilon_2}(\tau)\right] + \left[\beta_G + \beta_I(\mu_x + Q_{\epsilon_1}(\tau))\right]g_i$$
$$= \mu_\tau + \beta_\tau g_i\,,$$

where the quantile level $\tau \in (0,1)$ and $Q_Y(\tau|G_i = g_i)$ is the conditional quantile for quantile level $\tau$, and $\mu_\tau$, $\beta_\tau$ are the regression coefficients in quantile regression.

If there is no interaction effect in the initial linear model (i.e., $\beta_I = 0$), the coefficient $\beta_\tau$ will be a fixed constant which equals to the genetic main effect, i.e., $\beta_\tau = \beta_G$ for all $\tau \in (0,1)$. When GxE effect is present (i.e., $\beta_I \neq 0$), $\beta_\tau$ will vary across different quantile levels. Therefore, testing the differential $\beta_\tau$ across quantile levels $\tau$ provides evidence for GxE interaction at a given genetic locus without requiring the environmental factor E to be pre-specified.

### Estimation of quantile integral effect

If a SNP $G$ is a vQTL for trait $Y$, the slopes $\beta_\tau$ will differ in quantile regressions based on different quantile levels $\tau$.

$$Q_Y(\tau|G = g, \boldsymbol{C}) = \mu_\tau + g\beta_\tau + \boldsymbol{C}\alpha_\tau$$

Here, $\boldsymbol{C}$ and $\alpha_\tau$ denote $m$ covariates and their regression coefficients. $\mu_\tau$ is the intercept. Let $\theta_\tau = (\mu_\tau, \beta_\tau, \alpha_\tau)$ denote the parameters in the quantile regression. These parameters can be estimated by minimizing the loss

$$\hat{\theta}_\tau = argmin_{\theta_\tau} \sum_{i=1}^{n} \rho_\tau(Y_i - \mu_\tau - g_i\beta_\tau - \boldsymbol{C}_i\alpha_\tau)$$

where $\rho_\tau(u) = u[\tau - I(u < 0)]$ is the quantile regression loss function and $i$ is the index for the $i$-th individual in the analysis. As demonstrated in the main text, estimating $\hat{\beta}_\tau$ by fitting quantile regression for a grid of $\tau$ levels for each SNP is computationally challenging in genome-wide analysis. Therefore, we developed a rank-score-based method to estimate $\hat{\beta}_\tau$ without the slow optimization process.

We show that the squared root of the rank-score test statistic $T_\tau$ is highly consistent with the squared root of the Wald test statistic $W_\tau$ (**Supplementary Figure 7**), i.e.,

$$\sqrt{W_\tau} = \frac{\hat{\beta}_\tau}{SE(\hat{\beta}_\tau)} \approx \sqrt{T_\tau} = \frac{S_\tau}{SE(S_\tau)}.$$

Here, $SE(\cdot)$ is the standard error and $S_\tau$ is defined as

$$S_\tau = \frac{1}{\sqrt{n}} \sum_{i=1}^{n} \hat{a}_i(\tau) G_i^*,$$

where $\hat{a}_i(\tau) = \tau - I(Y_i < \boldsymbol{C_i}\hat{\alpha}_\tau)$, $\hat{\alpha}_\tau$ is the estimated coefficient under the null model, i.e.,

$$\hat{\alpha}_\tau = argmin_{\alpha_\tau} \sum_{i=1}^{n} \rho_\tau(Y_i - \boldsymbol{C_i}\alpha_\tau),$$

$I(Y_i < \boldsymbol{C_i}\hat{\alpha}_\tau)$ is a binary indicator for whether $Y_i$ is smaller than the estimated $\tau^{th}$ conditional quantile for $Y_i$, $\boldsymbol{C} = [C_1, \ldots, C_n]^T$ is the $n \times m$ matrix for covariates, $G_i^*$ is $i$-th element of $G^* = (I - P_{\boldsymbol{C}})G$ where $P_{\boldsymbol{C}} = \boldsymbol{C}(\boldsymbol{C}^T\boldsymbol{C})^{-1}\boldsymbol{C^T}$ is the projection matrix onto the column space of $\boldsymbol{C}$, and $G = [G_1, \ldots, G_n]^T$ is the $n$ dimensional genotype vector. Hence, we have

$$\hat{\beta}_\tau \approx \frac{S_\tau SE(\hat{\beta}_\tau)}{SE(S_\tau)}.$$

Since the calculation of $S_\tau$ and $SE(S_\tau)$ does not require minimizing the loss, what remains is to find a proxy for $SE(\hat{\beta}_\tau)$ without the fitting process. We consider the following two quantile regressions

$$Q_Y(\tau|G, \boldsymbol{C}) = \mu_\tau + G\beta_\tau + \boldsymbol{C}\alpha_\tau,$$

and

$$Q_Y(\tau|d, \boldsymbol{C}) = b_\tau + d\gamma_\tau + \boldsymbol{C}\alpha_\tau,$$

where $d \sim N(0,1)$ is independent with $\boldsymbol{C}$. The ratio of the kernel estimator[1] of $SE(\hat{\beta}_\tau)$ and $SE(\hat{\gamma}_\tau)$ is

$$\frac{SE(\hat{\beta}_\tau)}{SE(\hat{\gamma}_\tau)} = \frac{\sum_i^n I(|Y_i - \boldsymbol{C_i}\hat{\alpha}_\tau - G_i\hat{\beta}_\tau| < c_n)}{\sum_i^n I(|Y_i - \boldsymbol{C_i}\hat{\alpha}_\tau - d_i\hat{\gamma}_\tau| < c_n)},$$

where $\boldsymbol{C_i}$ is the i-th row of the covariates matrix $\boldsymbol{C}$, $G_i$ is the i-th element of the genotype vector $G$, $d_i$ is the i-th element of the vector $d$, $c_n$ is a quantity that is same for denominator and numerator and $I(.)$ is the indicator function. We note that $d$ is a randomly simulated Gaussian variable and thus the effect size $\gamma_\tau$ is 0. In complex trait genetics, each SNP only has a weak effect on the phenotypic distribution[2]. Therefore, effect size $\beta_\tau$ is also close to 0 and the following ratio is approximately equal to 1

$$\frac{SE(\hat{\beta}_\tau)}{SE(\hat{\gamma}_\tau)} \approx \frac{\sum_i^n I(|Y_i - \boldsymbol{C_i}\hat{\alpha}_\tau| < c_n)}{\sum_i^n I(|Y_i - \boldsymbol{C_i}\hat{\alpha}_\tau| < c_n)} = 1.$$

Therefore, we could obtain a computationally efficient rank-score estimate for $\beta_\tau$:

$$\hat{\beta}_\tau^{rs} = \frac{S_\tau SE(\hat{\gamma}_\tau)}{SE(S_\tau)}.$$

**Supplementary Figure 8** demonstrates the equivalence between a standard estimate $\hat{\beta}_\tau$ based on minimizing the loss and our rank-score estimate $\hat{\beta}_\tau^{rs}$.

Further, the explicit formula for $\hat{\beta}_\tau^{rs}$ is

$$\hat{\beta}_\tau^{rs} = \frac{S_\tau SE(\hat{\gamma}_\tau)}{SE(S_\tau)}$$
$$= \frac{n^{-\frac{1}{2}} \sum_{i=1}^{n} \hat{a}_i(\tau) G_i^* \, SE(\hat{\gamma}_\tau)}{n^{-1}\tau(1-\tau) {G^*}^T G^*}$$
$$= \frac{\sum_{i=1}^{n} Y_{i\tau} G_i^*}{{G^*}^T G^*}$$
$$= ({G^*}^T G^*)^{-1} {G^*}^T Y_\tau,$$
where $Y_\tau = [Y_{1\tau}, \dots, Y_{n\tau}]^T$ is a $n$ dimensional vector, and
$$Y_{i\tau} = \frac{\sqrt{n}\hat{a}_i(\tau) SE(\hat{\gamma}_\tau)}{\sqrt{\tau - \tau^2}}.$$
Then, the corresponding quantile integral effect with the rank-score estimate is
$$\hat{\beta}_{QI} = \frac{\sum_{k=1}^{K} \left(\hat{\beta}_{1-\tau_k} - \hat{\beta}_{\tau_k}\right)}{K}$$
$$= \frac{\sum_{k=1}^{K} \left[({G^*}^T G^*)^{-1} {G^*}^T Y_{1-\tau_k} - ({G^*}^T G^*)^{-1} {G^*}^T Y_{\tau_k}\right]}{K}$$
$$= ({G^*}^T G^*)^{-1} {G^*}^T \frac{\sum_{k=1}^{K} \left(Y_{1-\tau_k} - Y_{\tau_k}\right)}{K},$$
which is equal to the least squares estimate for the L2 loss
$$\hat{\beta}_{QI} = argmin_\beta \| Y_{QI} - {G^*}^T \beta \|^2$$
where $Y_{QI}$ denotes $\sum_{k=1}^{K} \left(Y_{1-\tau_k} - Y_{\tau_k}\right) / K$.

**QUAIL test statistic**

Consider the hypotheses
$$H_0: \beta_\tau = \beta_G, \text{for all } \tau \in (0,1),$$
$$H_1: \beta_\tau \text{ is not a constant},$$
where $\beta_G$ is a constant. We investigate the asymptotic distribution of $\hat{\beta}_{QI}$ under the null hypothesis.

Based on derivations shown in the last section, we know that
$$\hat{\beta}_{QI} = \left({G^*}^T G^*\right)^{-1} {G^*}^T Y_{QI} = \left({G^*}^T G^*\right)^{-1} {G^*}^T \frac{\sum_{k=1}^{K} \left(Y_{1-\tau_k} - Y_{\tau_k}\right)}{K}.$$
Given an arbitrary $\tau \in (0,1)$, we have
$${G^*}^T Y_\tau = \frac{{G^*}^T \hat{a}(\tau) \times SE(\hat{\gamma}_\tau)}{\sqrt{\tau - \tau^2}},$$
where $G^*$ is the $n \times 1$ vector of genotype residuals described above; $\hat{a}(\tau)$ is the $n \times 1$ vector of quantile rank scores for quantile level $\tau$ whose $i$-th element is $\hat{a}_i(\tau)$.

We assume the genetic effect on trait level (i.e., $\beta_G$) is small. Then, according to the rank-score inference[3],
$$E\left[{G^*}^T \hat{a}(\tau)\right] \to 0, n \to \infty \text{ for all } \tau \in (0,1).$$
When the assumption of small genetic effect on trait level (i.e., $\beta_G$) might violated, we first regress the phenotype $Y$ on genotype $G$ and use the phenotypic residual in this regression to perform analysis. This is to ensure the genetic effect on the trait residual level (i.e., $\beta_G$) is 0.

Based on the properties of the Powell sandwich standard error estimates[1],

$$SE(\hat{\gamma}_\tau) \to 0\,, n \to \infty \text{ for all } \tau \in (0,1).$$

Then, we have

$$E\left[G^{*T} Y_\tau\right] \to 0\,, n \to \infty \text{ for all } \tau \in (0,1),$$

and thus

$$E[G^{*T} \sum_{k=1}^{K} (Y_{1-\tau_k} - Y_{\tau_k})] \to 0\,,\ n \to \infty.$$

Since $\hat{\beta}_{QI}$ is the least squares estimator which is consistent and asymptotically normal. The limiting distribution of $\hat{\beta}_{QI}$ under the null hypothesis is

$$\hat{\beta}_{QI} \sim N\left(0, \sigma^2 \left(G^{*T} G^*\right)^{-1}\right),$$

where $\sigma^2 = Var(\epsilon)$ and $\epsilon$ is the residual in linear regression $Y_{QI} = G^*\beta + \epsilon$. Therefore, we can test the vQTL effect using $\hat{\beta}_{QI}$ and its limiting distribution. In our implementation, we use a linear regression $Y_{QI} = G^*\beta + \epsilon$ to obtain the QUAIL test statistic and p-values.

### QUAIL dispersion effect

In GWAS, we use a linear regression with SNP $G$ and covariates $\boldsymbol{C}$ to estimate the effect of SNP on the level of trait $Y$.

$$Y = \mu_\tau^{lm} + G\beta_\tau^{lm} + \boldsymbol{C}\alpha_\tau^{lm} + \epsilon^{lm}$$

Let $\hat{\beta}_\tau^{lm}$ denote the least squares estimate of $\beta_\tau^{lm}$. The estimate $\hat{\beta}_\tau^{lm}$ can be denoted in an equivalent form

$$\hat{\beta}_\tau^{lm} = \left(G^{*T} G^*\right)^{-1} G^{*T} Y^*,$$

where $Y^* = (I - P_{\boldsymbol{C}})Y$ and $G^* = (I - P_{\boldsymbol{C}})G$. $P_{\boldsymbol{C}} = \boldsymbol{C}(\boldsymbol{C}^T\boldsymbol{C})^{-1}\boldsymbol{C}^{\boldsymbol{T}}$ is the projection matrix onto the column space of $\boldsymbol{C}$. Define $Y_{disp}$ as

$$Y_{disp} = (I - P_{Y^*})Y_{QI},$$

where $P_{Y^*} = Y^*\left(Y^{*T} Y^*\right)^{-1} Y^{*T}$ is the projection matrix onto the column space of $Y^*$. Then, the estimated QUAIL dispersion effect is

$$\hat{\beta}_{QI}^{disp} = \left(G^{*T} G^*\right)^{-1} G^{*T} Y_{disp}$$

Note that $\mathrm{Cov}\left(Y_{disp}, Y^*\right) = 0$. We have

$$\begin{aligned}\mathrm{Cov}\left(\hat{\beta}_{QI}^{disp}, \hat{\beta}_\tau^{lm}\right) &= \mathrm{Cov}\left(\left(G^{*T} G^*\right)^{-1} G^{*T} Y_{disp}, \hat{\beta}_\tau^{lm} = \left(G^{*T} G^*\right)^{-1} G^{*T} Y^*\right) \\ &= \left(G^{*T} G^*\right)^{-1} G^{*T}\, \mathrm{Cov}\left(Y_{disp}, Y^*\right) G^* \left(G^{*T} G^*\right)^{-1} \\ &= 0\end{aligned}$$

**Supplementary Figures:**

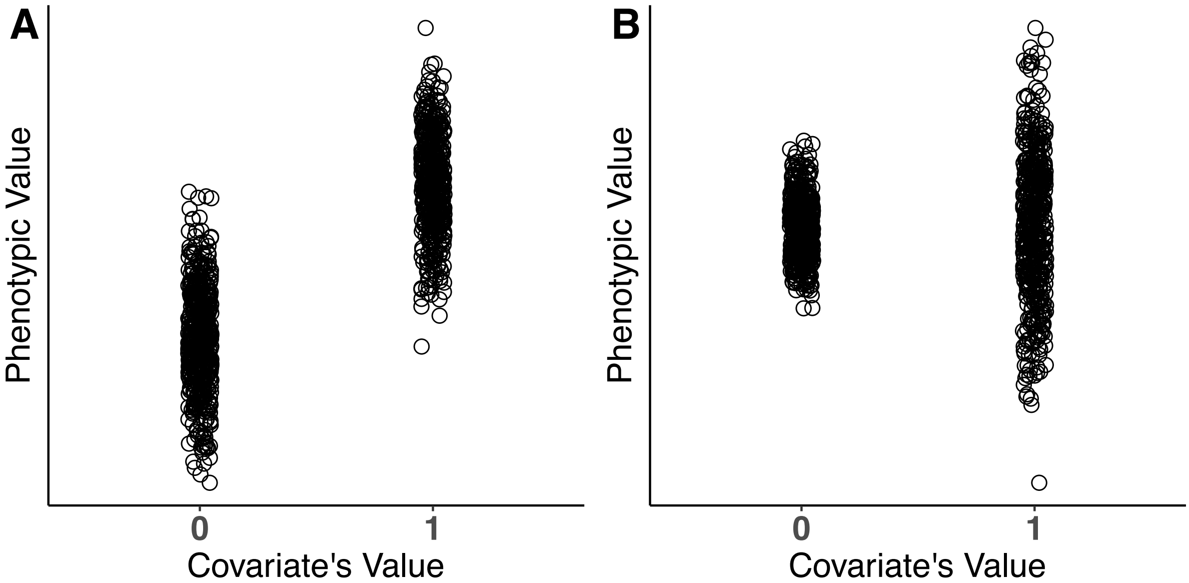


**Supplementary Figure 1. Covariates can affect the level and the variability of quantitative traits.** (**A**) This panel illustrates a scenario where the covariate only affects the phenotypic level. The phenotype is simulated as $Y = C\beta + \epsilon$, where $C$ is the covariate, $\beta$ is the covariate's effect, $\epsilon$ is the error term following $N(0,1)$. (**B**) Here, the covariate only has effects on phenotypic variance. The phenotype is simulated as $Y = \beta CE + \epsilon$, where $C$ is the covariate, $E$ is the environmental factor following $N(0,1)$, $\beta$ is the interaction effect, and $\epsilon$ is the error term following $N(0,1)$.

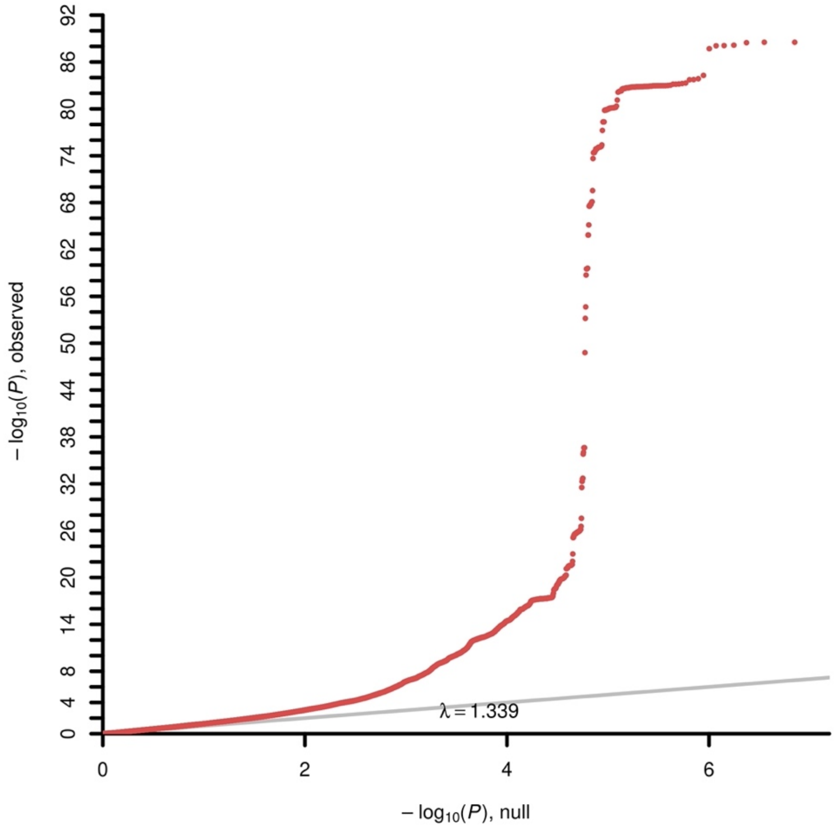


**Supplementary Figure 2. Quantile-quantile plot of QUAIL vQTL for BMI in UK Biobank.**

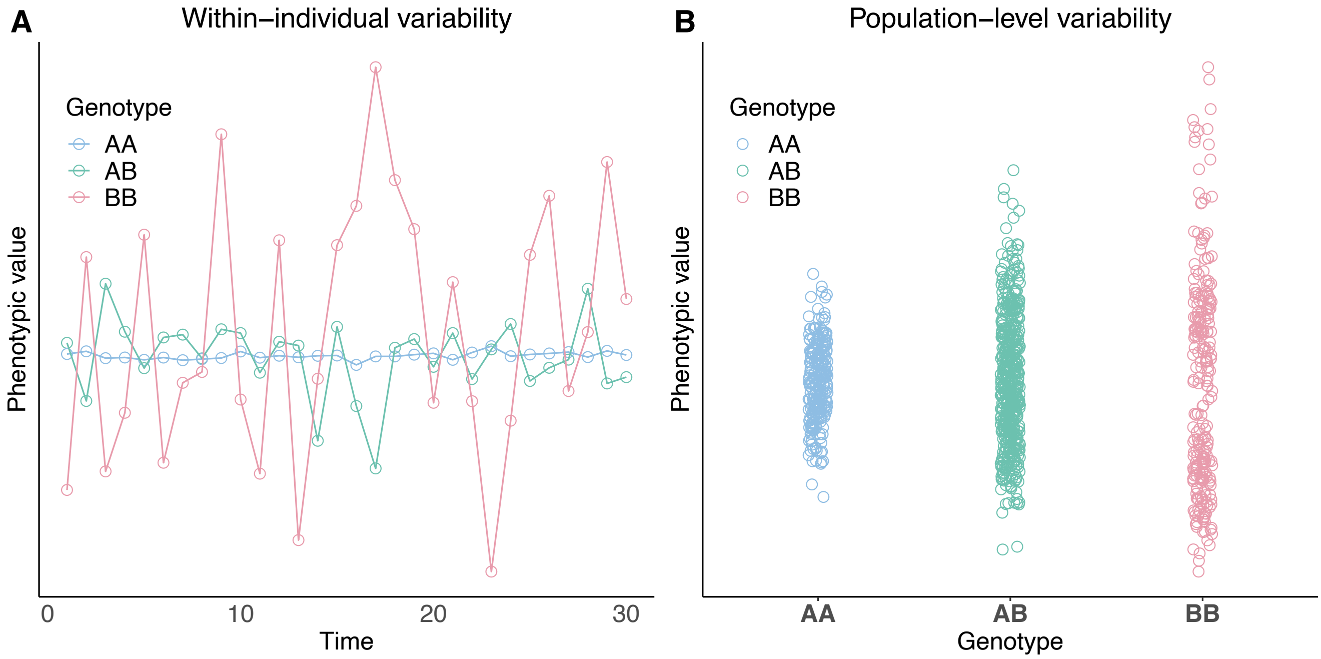


**Supplementary Figure 3. Genetic effects on within-individual variability and population-level variability.** (**A**) This panel illustrates the genetic association with within-individual trait variability. Individuals with more counts of allele B have increased phenotypic variability across time points. (**B**) This panel illustrates the genetic association with population-level trait variability. The subgroup of samples with the more counts of allele B has increased phenotypic variability between individuals.

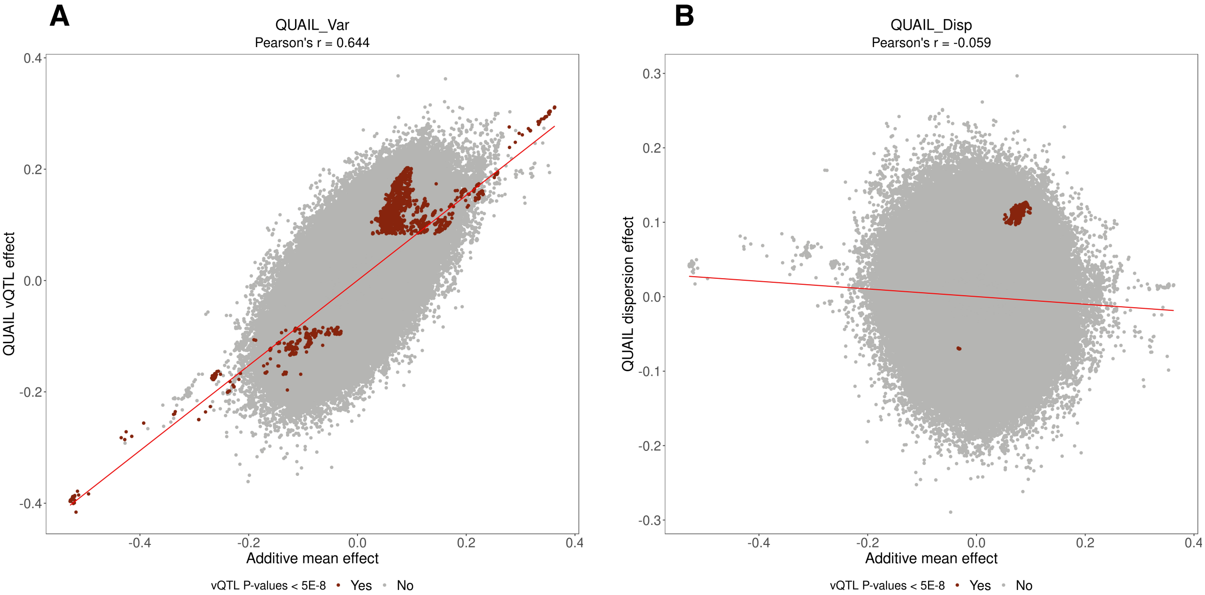


**Supplementary Figure 4. Relationship between the estimates for additive mean effect and variance effect.** The panels show results based on: (**A**) QUAIL, (**B**) QUAIL dispersion effect. Each data point is a SNP. X-axis indicates the estimated additive mean effect. Y-axis represents the estimated variance effect. The highlighted points represent the SNP with vQTL P-value < 5.0e-8.

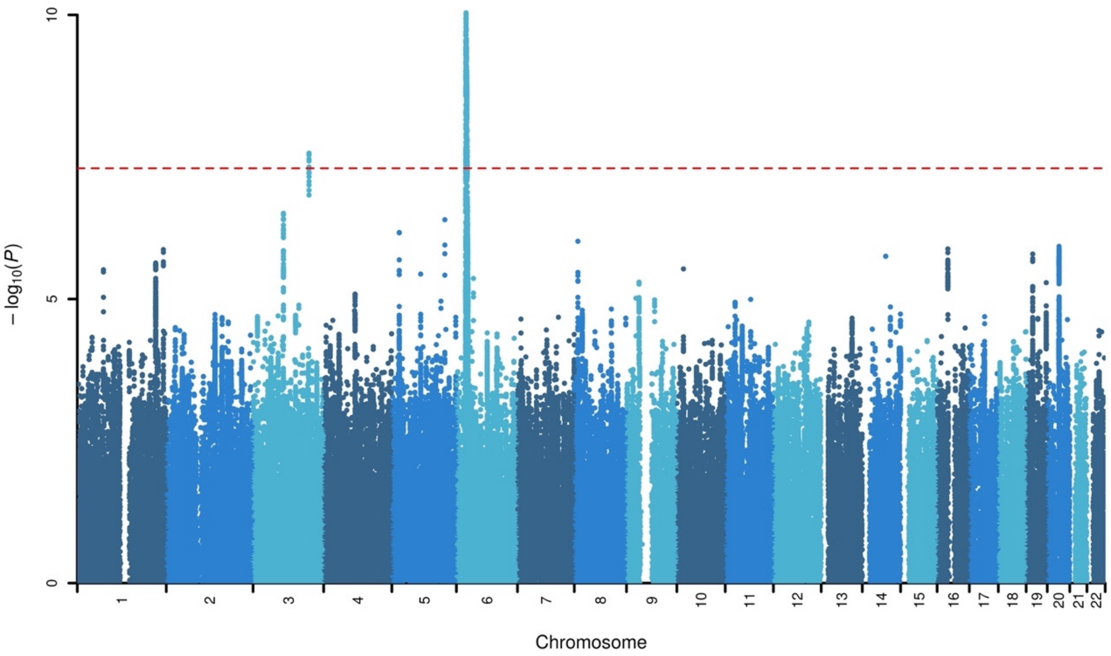


**Supplementary Figure 5. Manhattan plot of QUAIL dispersion effect for BMI in UK Biobank.** Dashed red line indicates Bonferroni-corrected significance threshold (P = 5.0e-8).

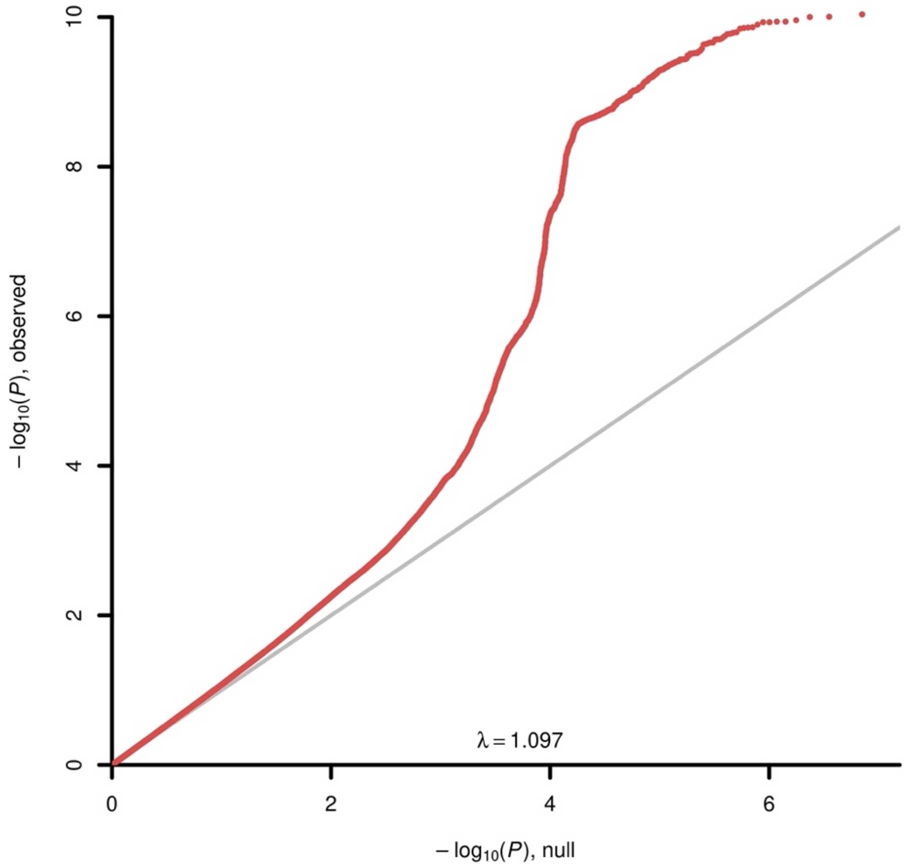


**Supplementary Figure 6. Quantile-quantile plot of QUAIL dispersion effect for BMI in UK Biobank.**

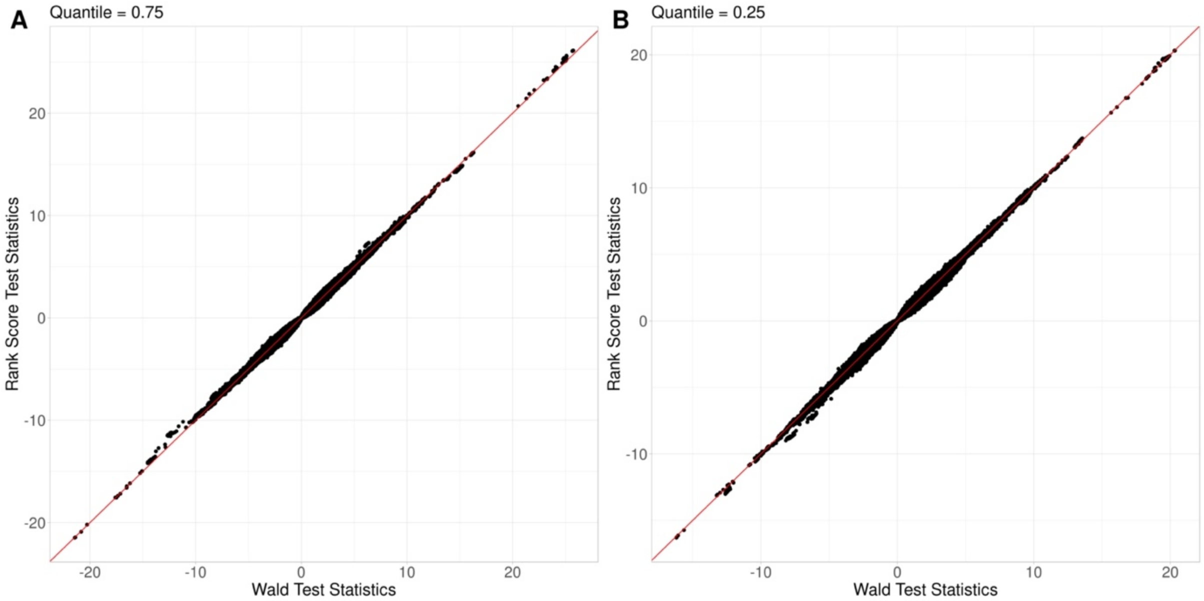


**Supplementary Figure 7. Comparison of Wald statistics and rank-score statistics.** Each point is a SNP. The X-axis is the Wald test statistics. Y-axis is the rank-score test statistics. **(A)** quantile level = 0.75 **(B)** quantile level = 0.25.

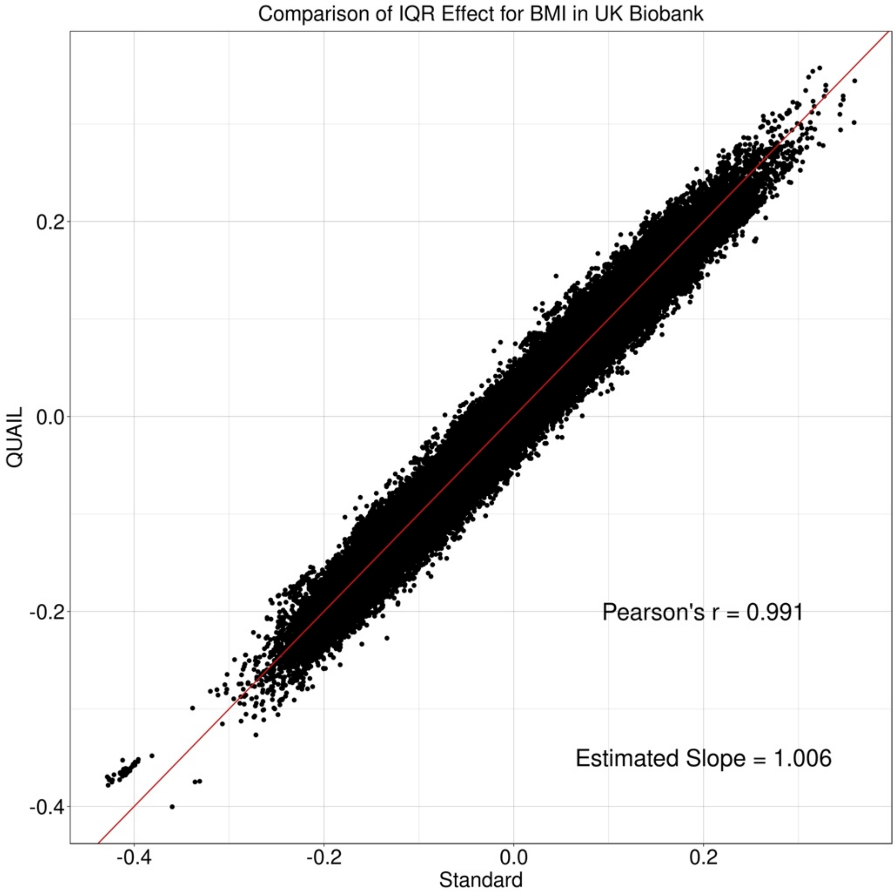


**Supplementary Figure 8. Comparison of effect sizes for interquartile range (IQR) given by QUAIL and standard quantile regression.** Consider a conditional quantile model of phenotype $Y$ given a SNP $G$ and a matrix $\boldsymbol{C}$ for $m$ covariates, $Q_Y(\tau|G,\boldsymbol{C}) = \alpha_{0\tau} + G\beta_\tau + \boldsymbol{C\alpha_\tau}$. The IQR effect is defined as $\beta_{0.75} - \beta_{0.25}$ from quantile regression. Each point is a SNP. The X-axis shows the estimated IQR effect obtained from the standard optimization approach. Y-axis shows the estimated IQR effect obtained from QUAIL.
